# Exploring and illustrating the mouse embryo: virtual objects to think and create with

**DOI:** 10.1101/2020.11.23.393991

**Authors:** Stefano Vianello

## Abstract

The teaching, learning, communication, and practice of Developmental Biology require interested parties to be at ease with the considerable spatial complexity of the embryo, and with its evolution over time as it undergoes morphogenesis. In practice, the four dimensionality of embryonic development (space and time) calls upon strong visual-spatial literacy and mental manipulation skills, generally expected to be innate or to come through experience. Yet it has been argued that Developmental Biology suffers the most from available traditional media of communication and representation. To date, few resources exist to engage with the embryo in its 3D and 4D aspects, to communicate such aspects in one’s work, and to facilitate their exploration in the absence of live observations. I here provide a collection of readily-usable volumetric models for all tissues and stages of mouse peri-implantation development as extracted from the eMouse Atlas Project (E5.0 to E9.0), as well as custom-made models of all pre-implantation stages (E0 to E4.0). These models have been converted to a commonly used 3D format (.stl), and are provided in ready-made files for digital exploration and illustration. Further provided is a step-by-step walkthrough on how to practically use these models for exploration and illustration using the free and open source 3D creation suite Blender. I finally outline possible further uses of these very models in outreach initiatives of varying levels, virtual and augmented reality applications, and 3D printing.

## Spatial thinking in Developmental Biology: understanding and communicating four dimensions

During mouse embryonic development a single, symmetric, undifferentiated cell transforms into a multicellular, differentiated organism populated by multiple specialised cell types (Pijuan-Sala et al., 2019). As new cells emerge, they migrate across the embryo, they squeeze between other tissues, they break down and deposit basement membranes, they traverse and integrate surrounding tissue layers (cfr. e.g. Arnold & Robertson (2009); Hashimoto & Nakatsuji (1989); Nahaboo & Migeotte (2018); Saykali et al. (2019); Viotti et al. (2014)). The embryo as a whole undergoes dramatic shape changes (McDole et al., 2018; Rivera-Pérez & Hadjantonakis, 2014; Snow, 1977). Students and researchers in mouse Developmental Biology are required to be familiar with such fundamental processes (gastrulation, cell migration out of the primitive streak, formation of the node, endoderm pocketing, gut morphogenesis, notochord deposition…). Yet, this requires them to build complex mental pictures involving simultaneous changes in multiple embryonic tissues, movement of different cell populations in three dimensions, and to entertain such complex mental representations as they evolve over time. Building such pictures is neither immediate nor intuitive, and for the many that do not have direct access to mouse embryos it usually only comes from the prolonged exposure to literature and the uncertain piecing together of a heterogeneous range of visual data formats. Here are then students and early career researchers scouring the literature for good representations of their structure of interest, at the right developmental timepoint, seen from that specific angle they need, with or without such and such other tissue. The outcome is rarely a single picture, and is typically a collage of disparate histological sections, immunostainings, electron microscopy images, diagrams.

Crucially to this point, the mental manipulation of 3D objects, over time (i.e. 4D), actively calls upon cognitive skills whose teaching has long been the focus of psychological and education research; notions of representational competence, spatial visualisation, spatial perception, visual-spatial literacy (see e.g. Milner-Bolotin & Nashon (2011); NRC (2012)). The importance of “the skill for thinking in three dimensions”, “for visualising shapes in the mind’s eye, rotating, translating, and shearing them”, and “imaging complex changes over time in the form of a cinematographic visual image” (Chadwick, 1978) will resonate strongly with most developmental biologists, just as that of so-called “penetrative thinking”, i.e. the capability to infer the cross-section of a layered object from surface features alone (Kali & Nir, 1996). While little research has focused on the importance of such skills in Developmental Biology Education (but see references in Hardin (2008)), foundational studies have been done in geoscience (and indeed compare the cross-section of a gastrulating embryo with the multi-layered landscape of earth crust sections). These studies have crucially shown that spatial thinking abilities can be i) taught, and ii) improved by practice (Kali & Nir, 1996; Titus & Horsman, 2009). Whether this transcends geological sciences or not, tools to foster visual-spatial literacy in Developmental Biology still remain scarce and not widely available (Hardin, 2008). It could be argued that the highly dynamic and dimensional nature of Embryology makes this discipline particularly penalised by traditional media of scientific dissemination.

When dealing with developmental data and with the inherent flatness (and stillness) of the printed page, the problem is not just the the extraction of information out of it (i.e. “understanding”), but also its deposition (i.e. “communicating”). Note the insidiousness of the problem: that is, even where one did successfully manage to mentally reconstruct complex representations of the system they want to illustrate, then its translation on paper still requires considerable artistic and technical ability. Regardless of the success of such translation process, it is this output that will serve as the basis for other scientists to inform their own mental pictures (cfr. Figure 1). The circle is vicious, and difficult to escape. A good illustration requires understanding, and a good understanding requires good summary references. If one does not have the possibility to see and explore real mouse embryos, dependence is built. Dependence on either luck and the pre-existence of illustrations of the embryonic context one wanted to illustrate, dependence on the pre-existing/solicited/paid work of rare scientists-artists, or dependence on the difficult investment of time and resources by the scientists themselves as to reach full illustrative autonomy.

**Fig. 1.**
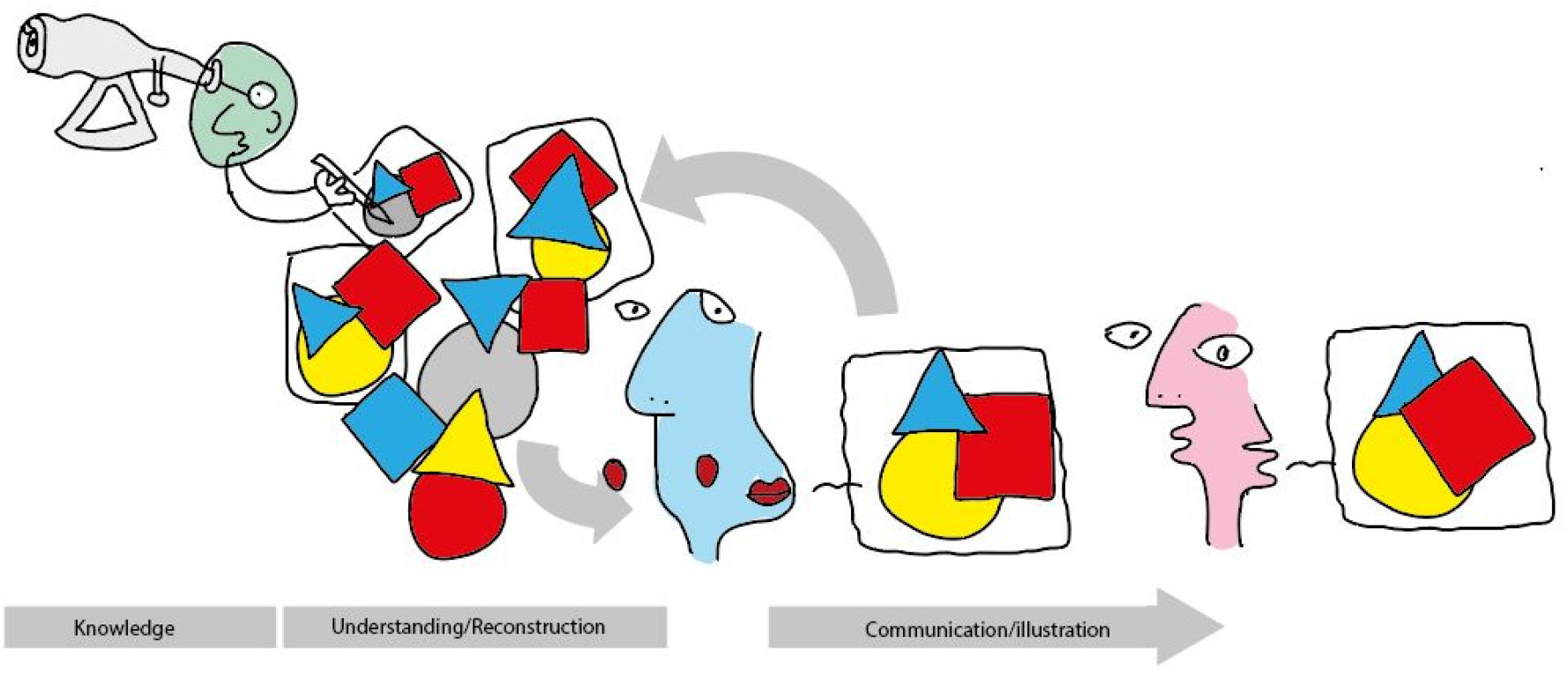
The vicious cycle of scientific illustration. Blue does not have access to real mouse embryos, and thus their entire understanding depends on the mental reconstruction of pre-existing representations from the literature. Blue’s output representations, regardless of the fidelity with which they match Blue’s mental reconstructions, will in turn serve as the basis of understanding for other scientist (e.g. Pink, and Blue themselves). Blue is thus entirely dependent on the output of direct observers such as Green, and has to wait for Green to draw the specific scenario needed.

How then to facilitate and democratise both the visualisation and the illustrations of mouse embryos? An important resource has been, and still remains, the Edinburgh Mouse Atlas Project (EMAP), the freely-accessible, annotated, online collection of 3D volumetric data hosted at https://www.emouseatlas.org/emap/ema/home.php, covering mouse development from pre-implantation, to gastrulation, to organogenesis and late post-implantation (Armit et al., 2017; Richardson et al., 2013). 3D models are easily navigable and allow users to define cutting planes as to see virtual histological sections at any desired angle. The value of such resource is maybe understated, but here are on-demand, user-explorable, customisable, 3D (+ timepoints) visualisations of mouse embryos at almost all stages of development. Clearly, this is a complete subversion of the normal illustration-spectator dependant relationship built by traditional developmental visualisation, and a powerful enabler of understanding (cfr. Figure 2). Crucial in this, is the role played by the actual data-format (i.e. a 3D object in space) and how it allows to completely evade the two-dimensional constraints of the printed page. Put very simply, an interactive, explorable 3D model condenses the informational potential of as many 2D pictures as there are angles to visualise that model.

**Fig. 2.**
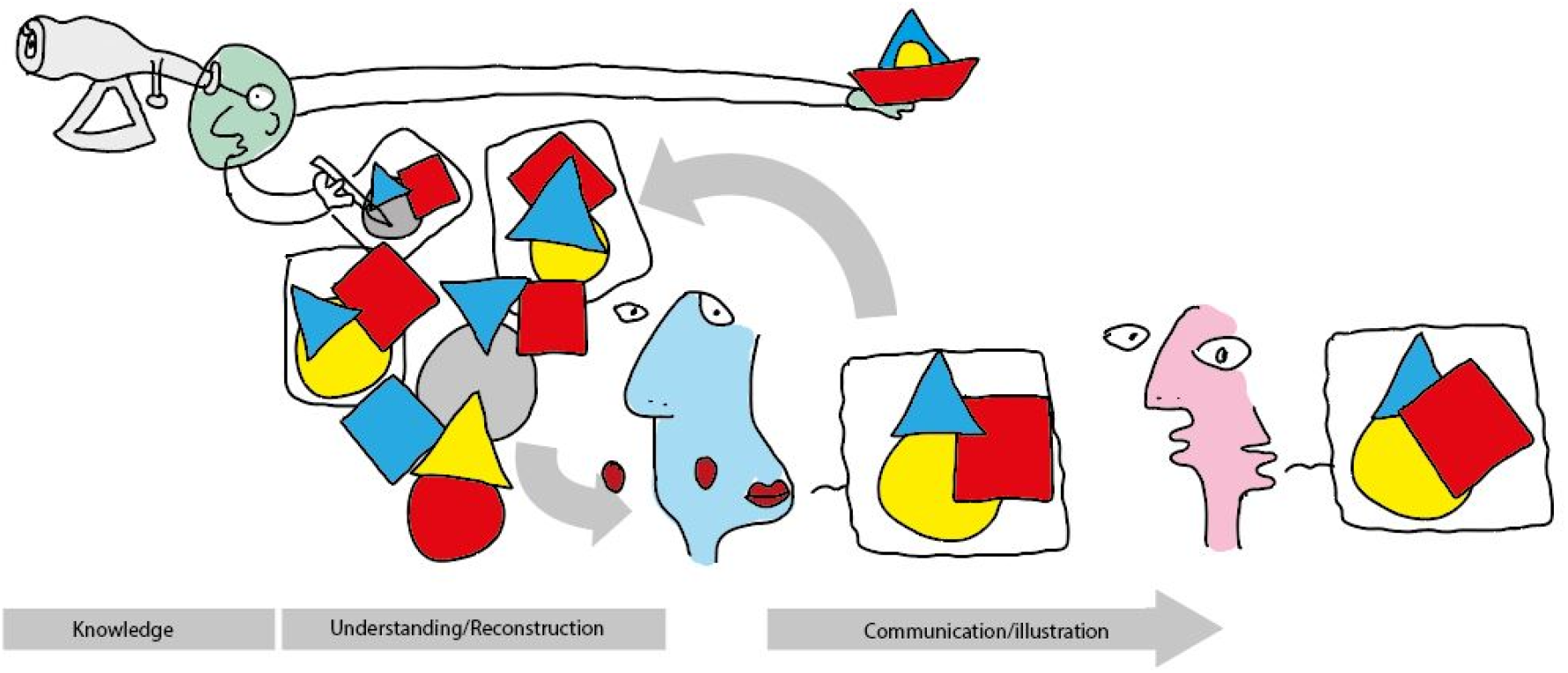
The power of 3D models. Direct observers like Green do not just illustrate the object from a specific point of view, but also decide to provide a 3D model of the original object (here, a boat). Blue can now observe the object from the specific angle they needed (becomes Green by proxy): the cycle of dependence is bypassed. Notice however that Blue cannot necessarily use the model provided by Green (e.g. only available as a video, or as 2.5D) when communicating with Pink. The cycle of dependence is broken at the understanding/reconstruction step, but not at the communication step.

With the evolution of our formats of scholarly communication, published data is also slowly starting to take life out of the page. Concomitant with the increasing use of experimental strategies outputting volumetric microscopy data, and with their application to the mouse embryo, such data is increasingly being presented as videos accompanying the online version of manuscripts. In these videos, the models are usually animated to rotate around their main axes, they might toggle specific structures on and off to reveal internal structure, or show internal cross-sections. Here is the dimensionality, here is the change over time, features so intimately associated with developmental biology (Hardin, 2008). The models live and are communicated in their 3D environment, if they are not user-explorable they are - at least - explored (cfr. Figure 2). Even if just provided in their printed version, 3D models still remain potent embryonic representations. One could say that the understanding of e.g. tailbud structure, embryonic heart development, and uterine architecture is almost immediate when shown, respectively, as in (Arora et al., 2016; Dias et al., 2020; Ivanovitch et al., 2017), even without the animated videos that may accompany such publications. Still, access to these models and thus the ability to explore them, is entirely dependent on the sharing practices of each individual set of authors, or of the venue of publication. Even in the case of the EMAP database, where volume data for every single embryonic structure is indeed downloadable (most updated version of the collection available at https://datashare.is.ed.ac.uk/handle/10283/2805), this does not come in a format of easy transfer to common visualisation and illustration softwares. These conversion steps, and the digital reuse of 3D objects, requires skills that - while not difficult - are yet not mainstream compared e.g. to the relative familiarity one is nowadays expected to have for 2D illustration software.

In the light of the “illustration -> understanding -> new illustration” framework outlined above, we finally also have to reflect on the usability of our visualisations. That is, to what extent do they allow the reader to transition from passive “visualiser” and “learner”, to active user, “illustrator”, “teacher”, “communicator”? To what extent do they empower them? To what extent to they break this cycle of dependency? Is it possible to imagine a framework where illustrations not only catalyse understanding, but they simultaneously facilitate creative experimentation? Where the lifecycle of an illustration/model does not end upon publication but gets new life in the hands of the reader? There is an untapped value in 3D models, probably uniquely so for the mouse developmental biology community given the existence of repositories such as EMAP. Tapping into it only requires easy access to the models themselves and the technical know-how on how to use and explore such models digitally. And here are scientist that will be able to explore these models on their own. Scientists and students that will be able to incorporate them into their own illustrations. Outreach officers that will be able to deploy them in e.g. virtual or augmented reality applications. 3D printers that will be able to bring these structure to the hands of students and of the public (Figure 3).

**Fig. 3.**
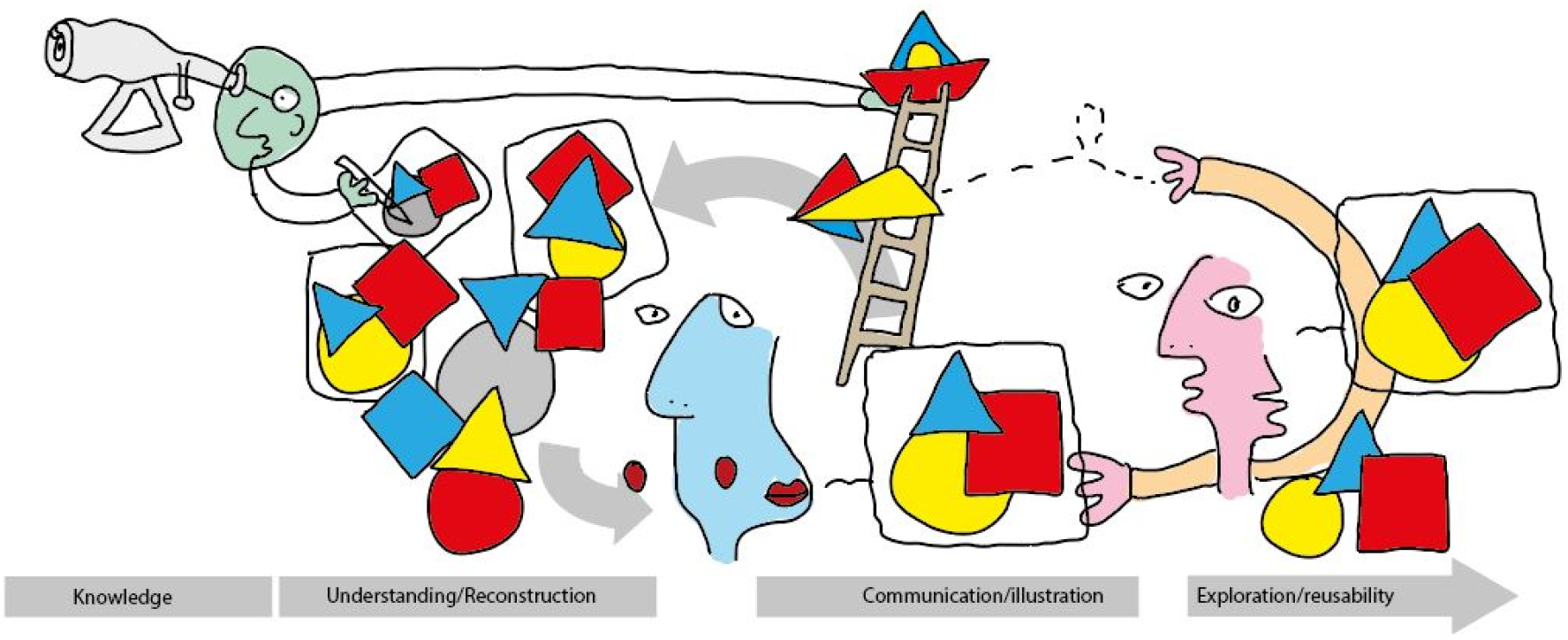
Visualisations become data. Green not only provides models, but makes them accessible too (e.g. copies of the model can be downloaded and manipulated). Blue can actively explore the object and is even more likely to find the point of view they needed. Pink developed the technical skills to use the model itself, and can now transform/reinterpret it, 3D print it, or use it in new visualisations. All of these elements feed back into the visual data formats available to the community, and thus further catalyse understanding.

I here provide a collection of readily-usable volumetric models for all tissues and stages of mouse peri-implantation development as extracted from the eMouse Atlas Project (E5.0 to E9.0), as well as custom-made models of all pre-implantation stages (E0 to E4.0). These models have been converted to a commonly used 3D format (.stl), and are provided in ready-made files for digital exploration and illustration. I further provide a step-by-step walkthrough on how to practically use these models using the free and open source 3D creation suite Blender. I finally outline possible further uses of these very models in outreach initiatives of varying levels, virtual and augmented reality applications, and 3D printing. I hope these resources and instructions may add to those already available to the mouse developmental biology community, serve as a soft introduction to more members of our community to the world of 3D illustration, and encourage creative exploration and experimentation in such a visually rich and stimulating field that is that of embryology.

### A collection of volumetric models of embryonic stages

An overview of all 16 models available for download is provided in Figure 4. These models cover Theiler Stage (TS) 01 to TS14, and include 8 custom-made models (covering TS01 to TS06, or E0.5 to E4.5), as well as 8 models assembled from EMAP data (covering TS07 to TS14, or E5.0 to E9.0). Also provided are two additional models corresponding to cross-sectional views of the early and late blastocyst models (TS04 and TS05, or E3.0 and E4.0), as to reveal the inner cell mass. As each of the 18 embryonic models provided is given as an assembly of individual subcomponents, a total of 251 individual volumes are thus available for download, exploration, and reuse (see Supplementary Data Table for a complete list).

**Fig. 4.**
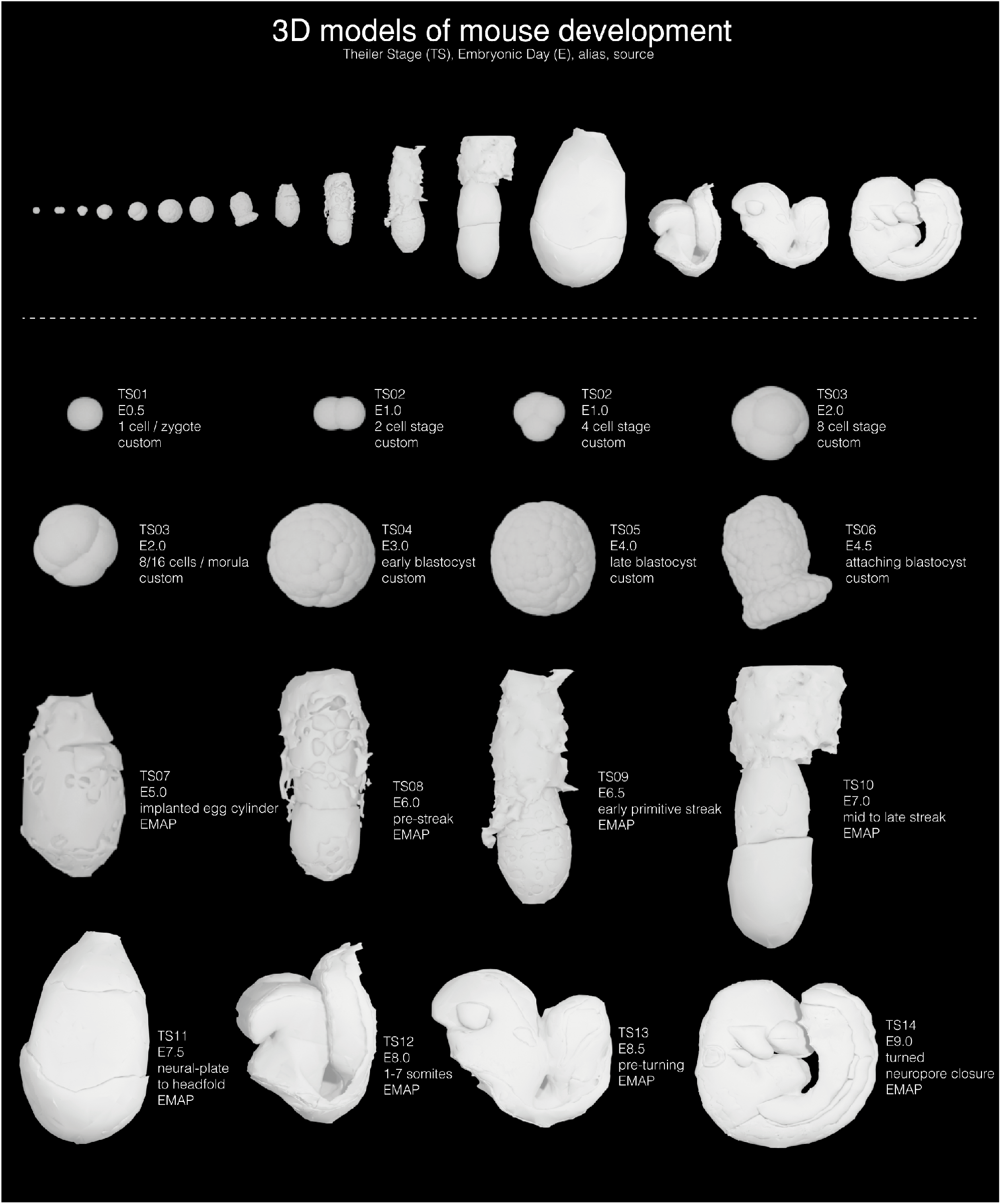
Summary of the 18 assembled models available for download. for a total of 251 individual volume models. Models for early and late blastocysts are also given in an open version showing the inner cell mass (not shown here).

The models have been deposited at http://doi.org/10.5281/zenodo.4284380 and will be found as 18 separate .blend files (one for each of the 16 stages, +2 open blastocysts). These files are to be opened with the free and open source 3D creation suite Blender (https://www.blender.org/) and have been prepared as to be immediately usable for exploration and illustration (see later paragraphs). Also provided are the 251 separate volumes of each embryonic structure represented, grouped by embryonic stage. These are provided as .stl format, as to be readily used not only for illustration, but also in 3D printing, gaming, and virtual/augmented reality environments. Finally, an additional .blend file with the full embryonic collection is also provided (as shown in Figure 4), as well as a “starting-pack” of pre-made materials to apply to the models (see Figure 5, and **Glossary**).

**Fig. 5.**
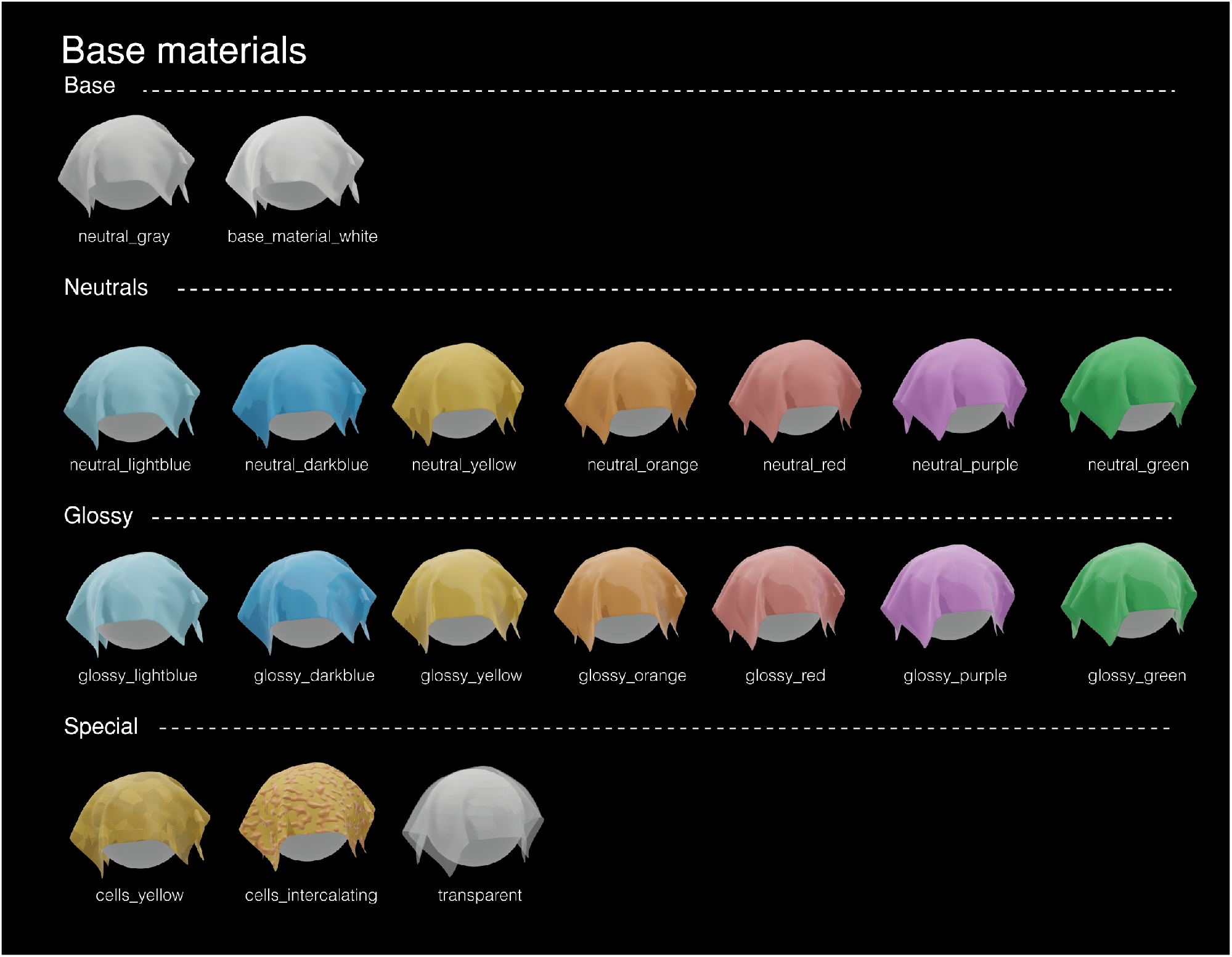
List of pre-made materials available for download. By default, 3D models will be displayed as grayish-white, but the visual appearance of a model can be controlled by applying materials/shaders to it. To facilitate the process, the above materials can be downloaded and used. These correspond to all the main colours, in a neutral and in a glossy (“wet”) version. “Special” materials include one made to mimic the surface of an epithelium, one showing mesenchymal cells migrating above it, and one making the structure transparent. Technical details on how these materials were created are provided in the **Methods section**.

### Use of the models for exploration/understanding

As mentioned in the introduction, one of the biggest advantages of having access to 3D models is that they allow the user to take control over the exploration of these very models (as per the paradigm illustrated in Figure 3). A big part of the models provided here have been long accessible and explorable on the EMAP portal, but they come in a dedicated file format to be open and read with the Java-based viewer JAtlasViewer. Through this viewer, the user can relate 3D structure and histological cross-sections at any cutting plane defined in the application. At the cost of losing the possibility to view cross-section histology (which would however still be available on the EMAP website), Blender provides an alternative way of manipulating and exploring 3D objects, while also allowing easy customisation of the models, rendering to 2.5D illustrations, the creation of scenes with multiple models, sculpting, and more advanced functionalities. Furthermore, and for 3D models that do not benefit from curated interactive visualisation platforms such as those of EMAP, Blender simply provides an easy-to use interaction and exploration platform.

To explore the embryonic models provided, just download the corresponding .blend file. If Blender has been installed on the computer, the file should then be openable just by double-clicking on it (Figure 6.1, here TS13 as an example). One can now freely explore the embryo. Unwanted tissues and structures (e.g. outer layers hiding internal structures) can be toggled off by clicking on the eye icon next to their name in the lateral menu (Figure 6.2). If one is unsure of the name of the tissue to be removed, right clicking on it will highlight its name in orange in the lateral list (later stage models are made up of up to 80 components!). As a rule of thumb, one will likely want to toggle off all extraembryonic tissues: these can be easily identified because their name is preceded by an “X” (see Supplementary Data Table). In the example provided here, most extraembryonic tissues have been removed, as well as mesenchmyal and cardiovascular structures around the tailbud to get a better view of gut tube and notochord.

**Fig. 6.**
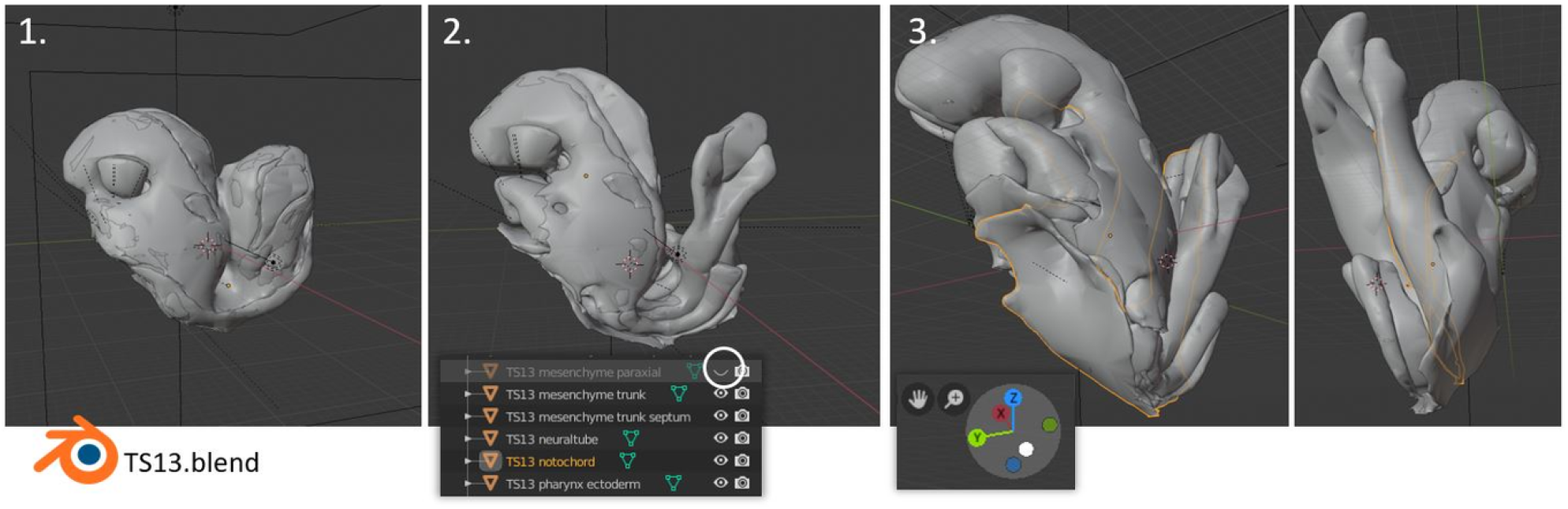
Exploring 3D embryonic models in Blender: moving perspective.

Exploration of the model can then proceed by scrolling in/out with the centerwheel of the mouse (zoom in/out), moving the mouse while holding the wheel down (rotation around the model), and moving the mouse while holding SHIFT and the wheel down (panning vertically and horizontally). Alternatively, and from Blender 2.8 onward, one can more simply interact with either of the three icons on the top right of the viewer (see Figure 6.3). In the example, this has been done to get a better view of how the endoderm pockets into the anterior of the embryo (left), and to track the notochord as it emerges from the tailbud (right).

An alternative way to explore the embryo models provided is to move and rotate the model itself, and to move structures in and out of them (see Figure 7, TS09 as an example). To move the whole embryo at once, select the “HANDLE” object in the list on the right (left-click on the name, Figure 7, bottom). The model can be now moved along the three axes, or rotated around them, by clicking on the corresponding icons at the left of the screen and then interacting with the widgets that appear on top of the model (Figure 7.2, move icon; Figure 7.3, rotate icon). The same actions can be performed with the keyboard shortcuts G (for “grab”, followed by X, Y, or Z to lock the movement along these axes), and R (for “rotate”, followed by X, Y, or Z to lock the rotation around these axes). The same can be done on individual components, rather than on the whole embryo, by selecting them first (right click on them in the viewer, or left click on their name on the right panel). In Figure 7.4 this has been done on the (visceral) endoderm to move it down and reveal the epiblast and the primitive streak at one side of the embryo (some extraembryonic structures were also toggled off as explained before).

**Fig. 7.**
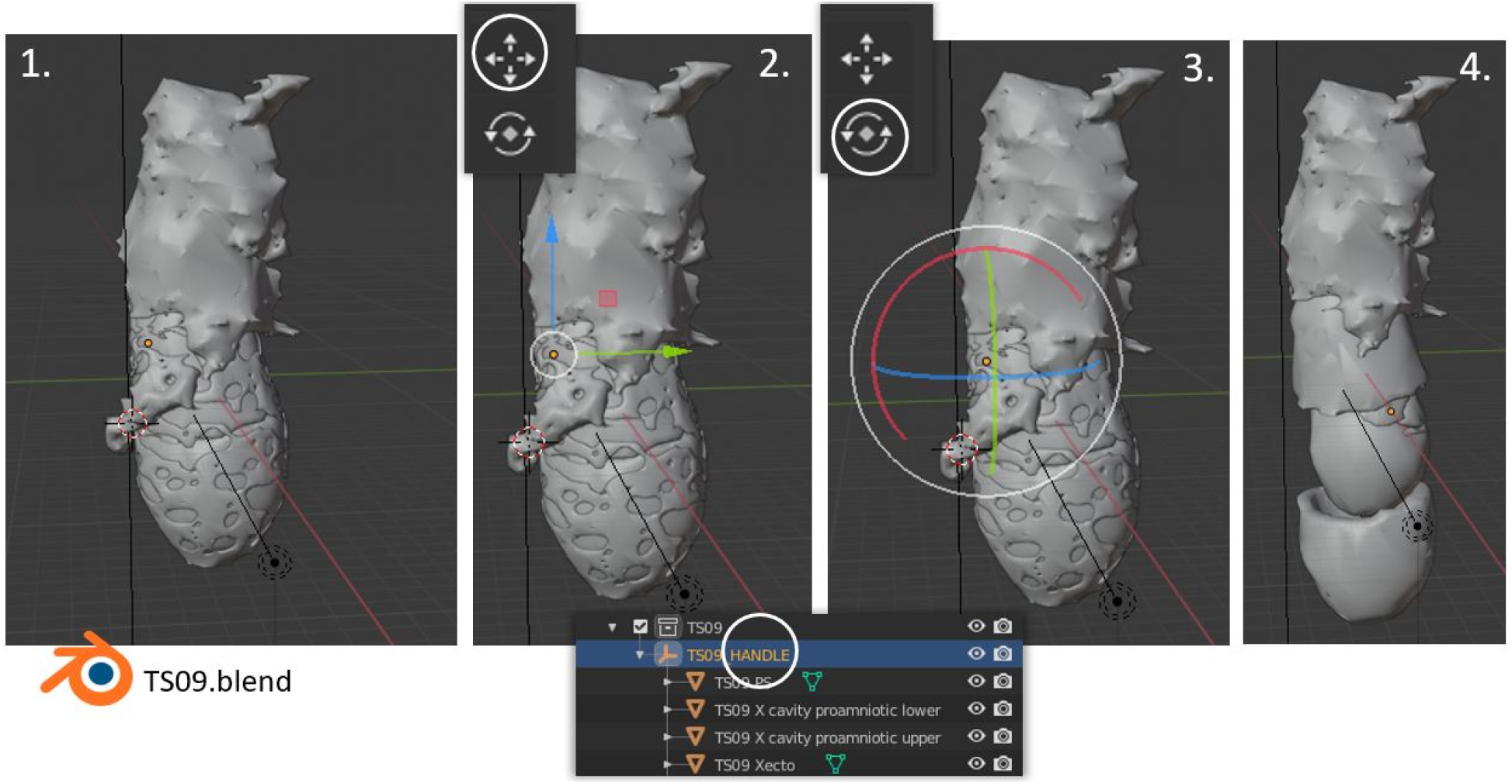
Exploring 3D embryonic models in Blender: moving the model.

Thus, the simple use of basic Blender functions (movement in space, on/off visibility toggling, object interaction and manipulation) allows for unbounded exploration of any of the embryonic models provided, allowing users to easily visualise and understand the spatial relationship of their structure of interest in the context of the whole embryo. In fact, it would be interesting to explore whether such exploration could improve visual-spatial literacy in the users, and whether this could represent a valid and beneficial tool within developmental biology pedagogical settings. To quote Hardin (2008): “the technology already exists to depict embryos on the computer as true 3D objects in 4D space. What is needed is the application of instructional materials development resources toward the production of such models. If such models become widely available, it should be possible to reclaim all four dimensions of the embryo in the undergraduate developmental biology curriculum”.

While these models have been here made available as pre-prepared .blend file, one can exploit these same basic functions to explore any type of volume data (.obj, .wrl, .stl; imported in a Blender scene via File>Import). This has been e.g. recently exploited for 3D exploration of cell migration tracking data in Samal et al. (2020), and can also be applied to explore .obj/.stl meshes generated from e.g. light-sheet imaging.

### Use of the models for illustration

The second advantage allowed by having access to 3D models in Blender is the possibility to create 2D (2.5D) illustrations out of them. This is done by taking snapshots of them at the angle needed: this creates an output image, a process called rendering (see **Glossary**), where the illustration software applies materials to the models (colour, roughness, refractivity…) and light to the scene. Regardless of drawing or artistic skills, one can thus generate reference figures for publications, scientific illustrations, material for outreach purposes, or even art. The illustration process thus becomes focused on the concept and message one wants to illustrate rather than on the technical aspects of it: accurate embryo models are already provided. Furthermore, illustrations of embryos from non-traditional perspectives do not require to mentally recalculate what the embryo would look like (think about how few illustrations show mouse embryos from the ventral or anterior side, compared to the lateral or dorsal sides).

To explain how the models provided here can be used for illustration, we will take a simple case study: that of a scientist wanting to illustrate endoderm development after gastrulation. This process involves extensive shape changes as the endoderm starts as a curved sheet on the surface of the embryo, later makes deep pockets at its two extremities, and finally closes to form a tube (the gut) while the whole embryo is closing on itself (Lewis & Tam, 2006). Illustration is made especially challenging in our case because the scientist is not confident they fully understand how the terminal endoderm pockets fit with other structures at the anterior and posterior of the embryo.

To create an illustration, one first need to set up a scene. In our specific case, the single-stage .blend files available for download are not sufficient: we would like to show models TS09 to TS14 within the same image. We will thus open TS09, and import the other models from the other files by selecting File>Append (Figure 8.1), navigating to the .blend file with the model needed (here TS10), opening its Collections folder, and selecting the TS10 entry. Appended models will be imported within the existing hierarchy, so make sure that you are in the topmost collection (Scene Collection, see Figure 8.1) before importing any model. Once the model has been imported and positioned (Figure 8.2; see previous section for how to move models in Blender), one can repeat the Append process to add each of the following embryonic stages (Figure 8.3). As explained before, specific structures and layers can be then toggled off (eye icon) to expose the structures one is interested in (here, the endoderm) (Figure 8.4).

**Fig. 8.**
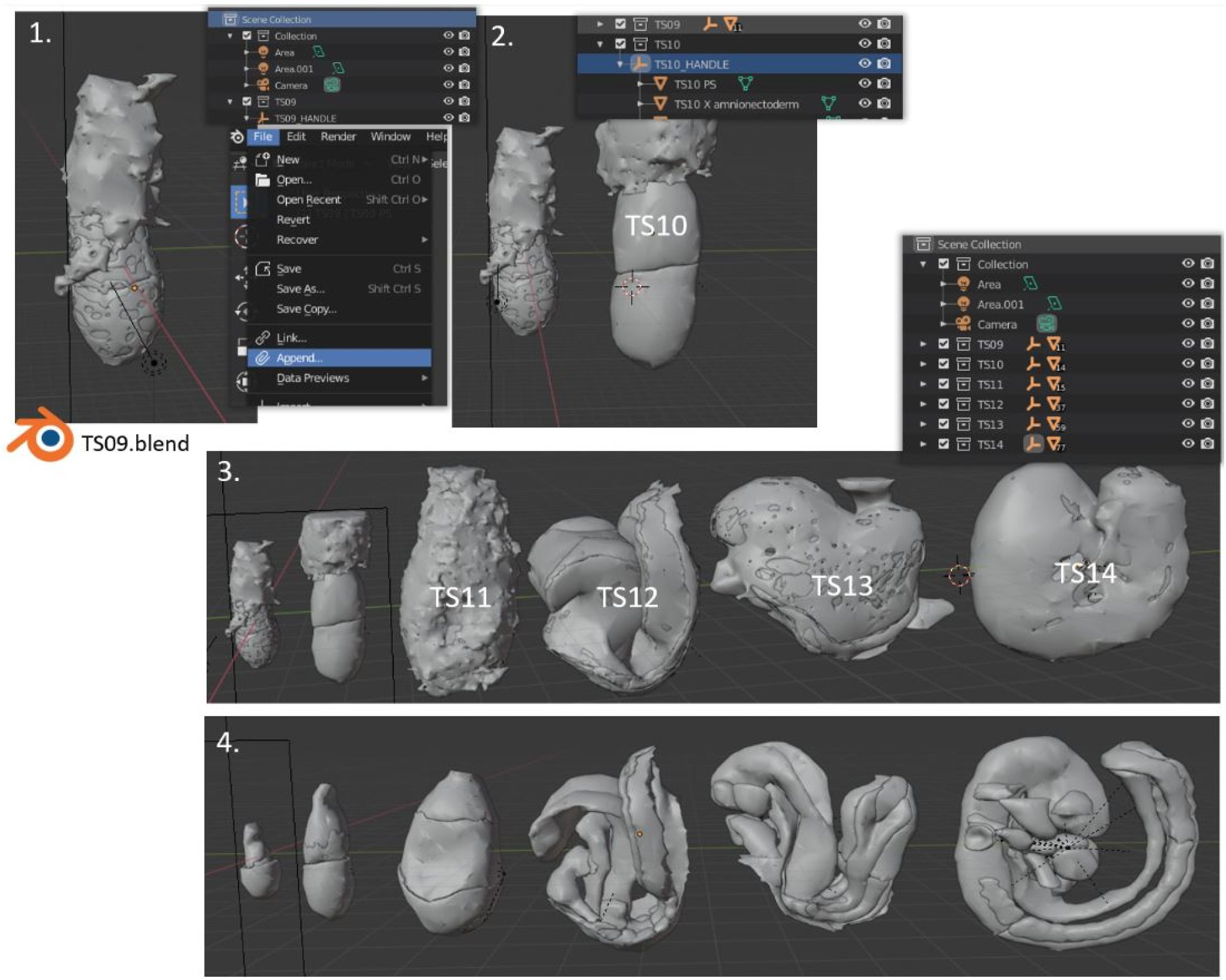
Importing models from other files by using File>Append>Collection.

Once the scene is set with all the models in the right place, it is time to add shaders/materials. Materials determine the final appearance (in its simplest form, the colour) of the models in the output illustration, and as such have a heavy influence on its success. To get an idea of what the final image will look like one can toggle the Rendered Viewport Shading icon at the top right of the viewport (Figure 9.2; allow some processing time). Since all models provided here come with a generic grayish shader, this is indeed how the models will look like (Figure 9.2). An important note is that, depending on where you placed your models into the scene, there might nor be enough light to actually see the real final colour of the model. This case is shown in Figure 9.3B, and corrected in Figure 9.3C by providing more light to the rightmost model. In Blender, light is extremely important, and every .blend model provided here comes with it’s own set of lamps. These are the objects highlighted in orange in Figure 9.3A, and can be imagined as literal rectangles shining light out of their surface: one illuminates the front of the model and the other the top. To add lights, select both of them (if using the name list on the right, Shift+LeftClick; if clicking directly on the objects, Shift+RightClick), press Shift+D to duplicate, and then move them to your next model (in Figure 9.2 you can indeed see how additional lights have been placed all along the lineup of models).

**Fig. 9.**
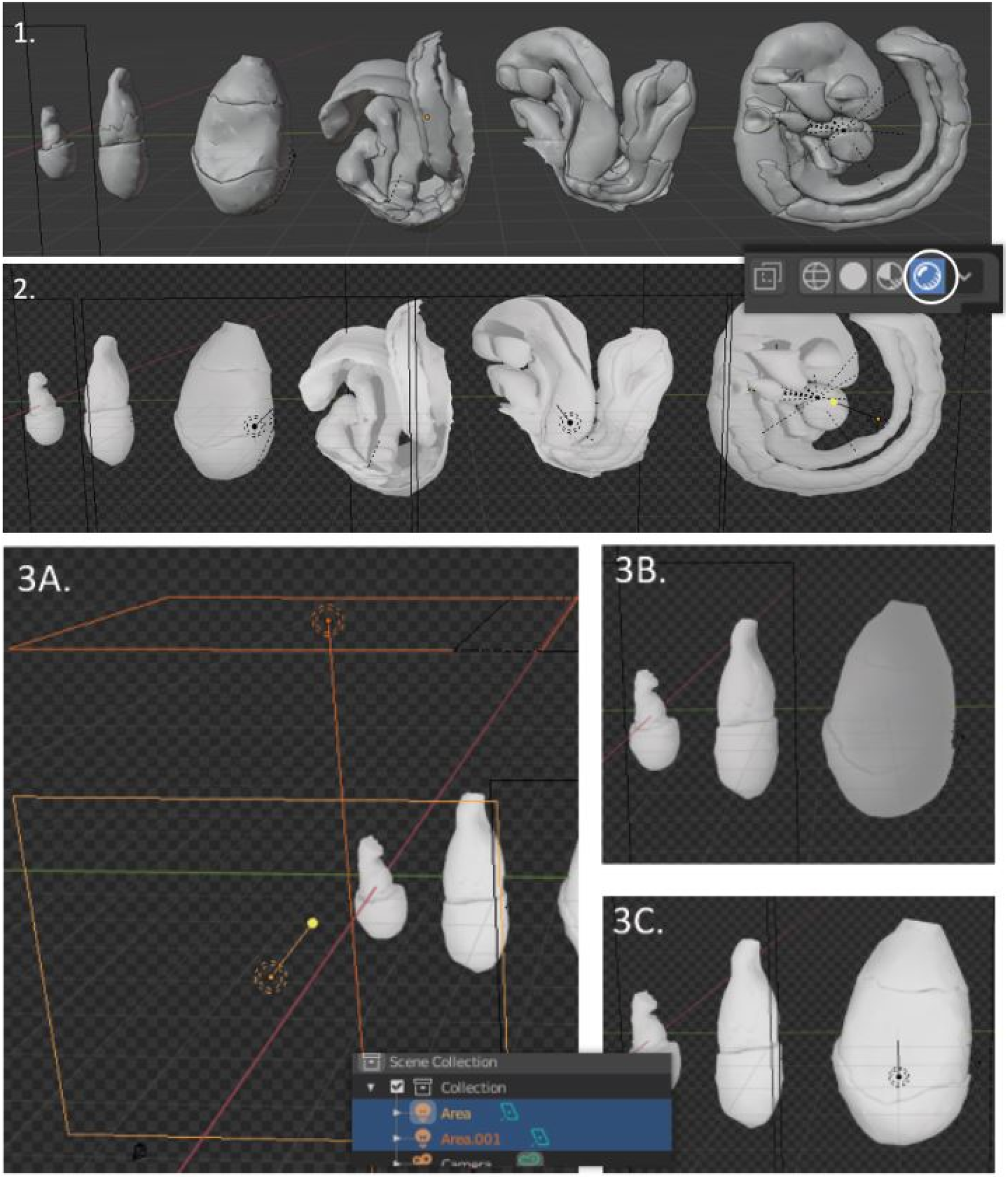
Rendered viewport and lights.

It’s then now time to apply materials to the structures we want to highlight (endoderm in this case). For a 2D illustration analogy, this is equivalent to having drawn a shape on paper and now having to colour it in. While Blender allows the creation of virtually any material one can imagine, we facilitated the task here by providing a starter-pack of pre-made ones (“materials.blend”, see Figure 5) alongside the embryo models. To import any of these materials into the scene, repeat the same process used to import additional embryo models (File>Append) but this time select the “materials.blend” file, and pick from the “Materials” folder. In this scene we will start by using the “cells_yellow” material. Once it has been imported, apply it by selecting the component you want to colour (here the visceral endoderm, Figure 10.1), clicking on the red checkered sphere icon in the panel on the right (this is the materials icon), and use the drop-down menu to select the material you just imported (Figure 11.1, right). Provided you are still in Rendered View mode (toggled on previously), the model will now look as in Figure 10.2. Proceed for all other models (materials only needed to be appended once) until satisfied (Figure 10.3). Since the endoderm soon starts being populated by cells intercalating from the mesoderm (Viotti et al., 2014), we have switched to the “cells_intercalating” material for all later models to better illustrate this phenomenon.

**Fig. 10.**
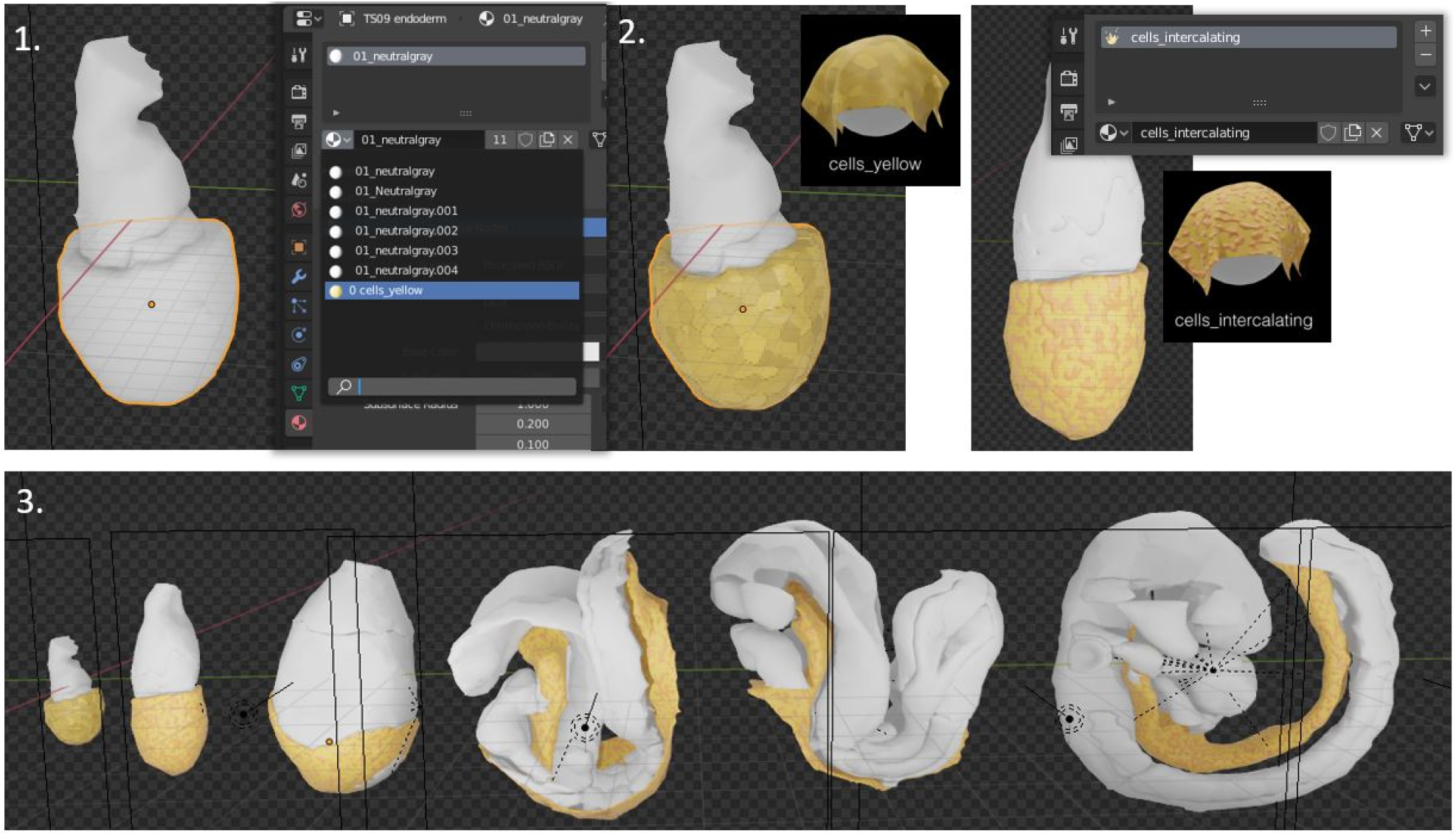
Applying materials.

**Fig. 11.**
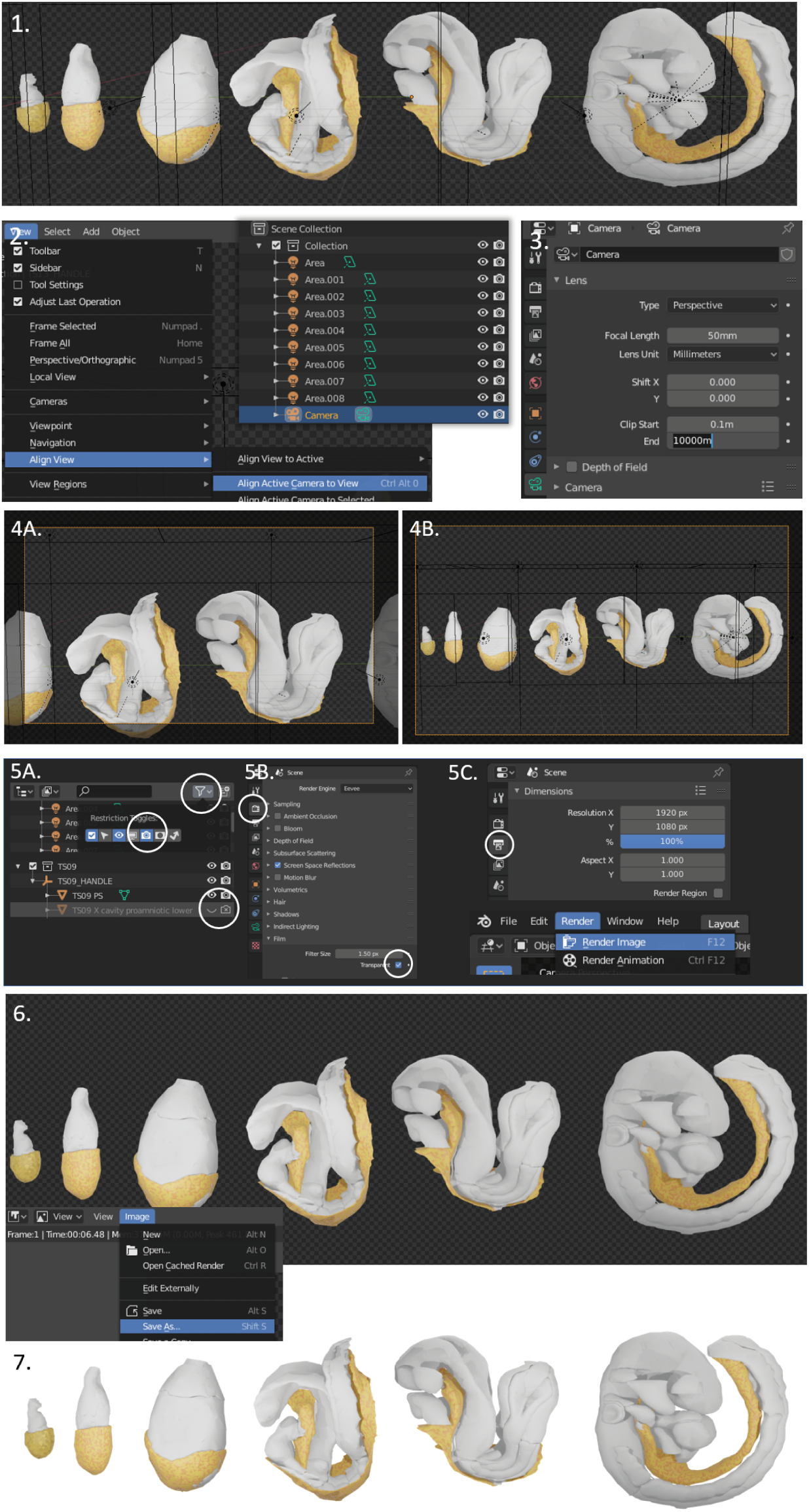
Camera positioning and rendering.

The models are in place, the scene is illuminated, and materials have been applied. We are now ready to transform this 3D scene into a 2D image. This is done by taking a virtual snapshot of our models, a process called rendering. This “snapshot” is taken with a Camera, which is indeed provided with all Blender scenes, just as lamps and lights are. To place the camera in position, start by positioning your view as to see your models as you want them in the final image (Figure 11.1). With the camera object selected (left-click on its name in the panel on the right), click on View>Align View>Align Active Camera to View (Figure 11.2): this will move the camera in position, and make you look through it. If all your models disappear at this point, this is because they happen to be out of the working distance range of the camera. To correct this, and with the camera object still selected, click on the green camera tab in the bottom right panel, and increase the End value until the models appear again (Figure 11.3, 10000 meters in this case). As in Figure 11.4A, the camera will not exactly capture the whole scene and will likely need to be moved “in” or “out” as to capture a smaller/bigger area. This is done by moving the camera along its relative Z axis, which is set through the keyboard shortcut “G” (for “grab”), followed by “Z” (lock movement to absolute Z axis), followed by “Z” again (lock movement to relative Z axis). Adjust the camera position until satisfied (Figure 11.4B): what you see through the camera is the snapshot that will be taken.

Once the camera is in place, it is time to launch the rendering process. To avoid any surprises, start by making sure that all the objects that had been hidden from view (eye icon toggled off), are also hidden from the final render. Just as you toggled off the eye icon next to the name of each object, toggle off the camera icon too (Figure 11.5A; if this icon is not shown, make it available from the Filters submenu as shown). Another aspect to double-check is that the “Transparent” Film checkbox is ticked in the display settings in the bottom left panel (tab with the camera icon, Figure 11.5B) because this will make the background transparent when the image is then saved as .png. Finally, the size/resolution of the final image can be set in the rendering tab (Figure 11.5C, tab with printer icon, here 1920px x 1080px). Notice that if you change the ratio of these values, your camera will also resize and you will need to realign it. Once all is set, launch the rendering via Render>Render Image (Figure 11.5C, bottom). A new window will open, and after some processing time, the final image will appear (Figure 11.6C). Save the image via Image>Save As. The image is done (Figure 11.7).

### Illustrating cross-sections

As a final illustration technique, a developmental biologist might often need to show cross-sections or cut-outs of a specific tissue: that is, not toggle the tissue off view completely, but just remove a section of it as to show underlying structures. This is achieved in Blender through the use of so-called Boolean modifiers, which essentially allow to substract/add shapes from/to one another. A cube or a parallelepiped can thus be intersected with our tissue of interest and the intersection deleted out. Upon removal of the cube, the tissue is left open: a process illustrated in Figure 12.

**Fig. 12.**
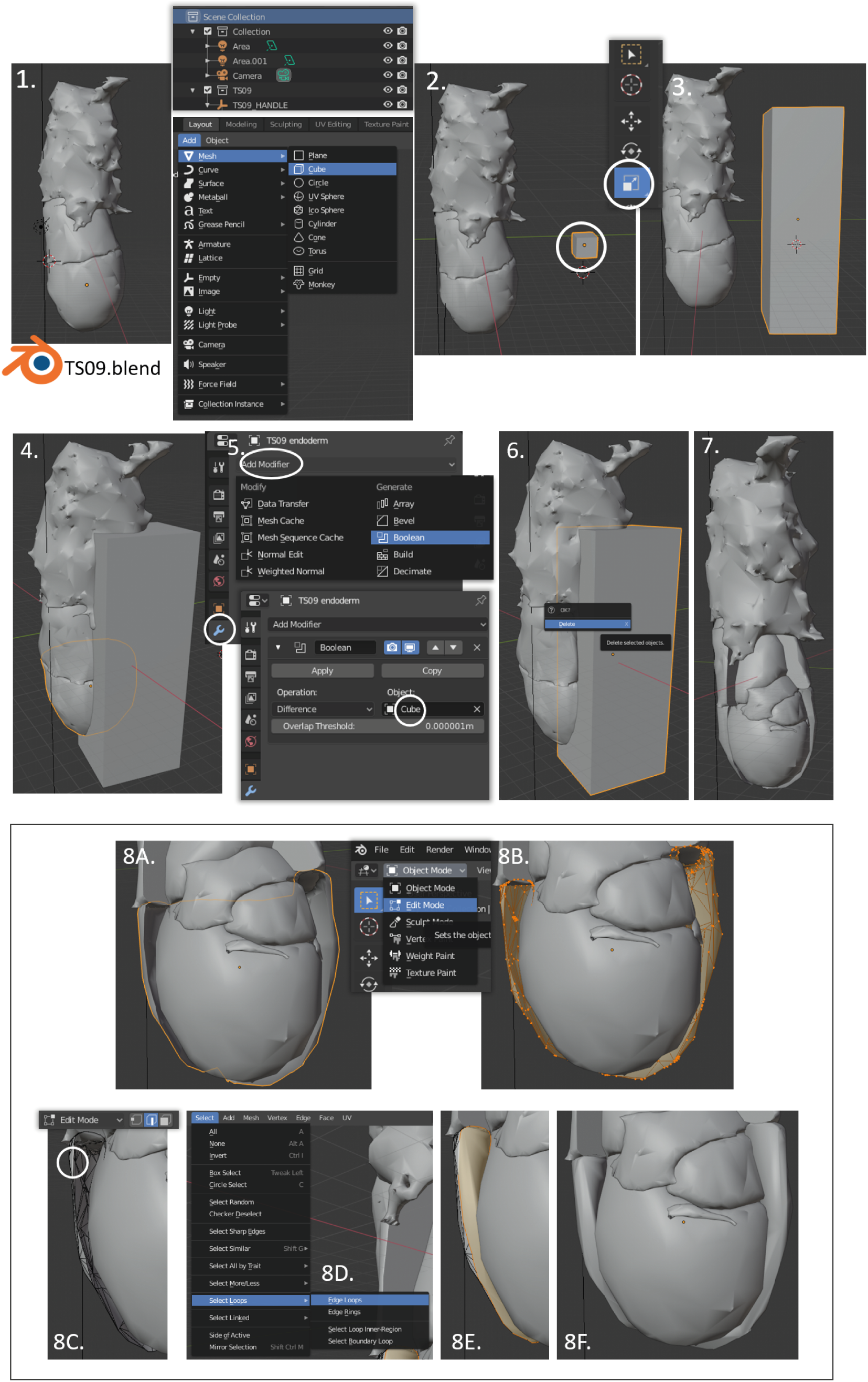
Creating cut-outs and using Edit Mode.

As a first step, one needs to create the shape to be subtracted. Making sure that one is in the topmost hierarchy (Scene Collection), add a cube to the scene by selecting Add>Mesh>Cube (keyboard shortcut Shift+A) (Figure 12.1). To scale the cube right-click on it (Figure 12.2) and select the Scale icon on the left, or use keyboard shortcut “S", followed by X, Y, or Z to lock the scaling to either of the three axes. Since we here want to cut out the visceral endoderm both in the embryonic and in the extraembryonic region, we make a parallelepiped that is as tall as the embryo (Figure 12.3). Once satisfied with the shape, move it as to intersect your model over the regions you want to carve out (Figure 12.4), select the tissue that needs to be cut (here starting with the embryonic visceral endoderm), and select the wrench icon of the bottom right menu (modifiers tab, Figure 12.5). Under “Add Modifier", select “Boolean”, and then make sure that the Operation is set to “Difference”, and the Object field displays the shape you want to subtract (our parallelepiped is called “Cube” in this example; Figure 12.5). Click on “Apply” and then repeat the whole process for the extraembryonic visceral endoderm. Once both tissues have been processed, delete the parallelepiped by pressing the keyboard shortcut “X” (Figure 12.6) to reveal what is left of the model (Figure 12.7). We can now clearly see epiblast and primitive streak under this newly created window through the visceral endoderm.

Because the models provided here are not “solid” (they are actually hollow, thin-layered envelopes), the kind of editing described above might reveal exposed holes in the cross-sections (Figure 12.8A, the cut visceral endoderm is hollow). These are problematic because one wants to hide this in the final illustration and give the impression of a solid model. To correct this, switch to Edit Mode (keyboard shortcut “Tab", or as in Figure 12.8A, right) to enable interaction with the mesh of the objects (Figure 12.8B). After deselecting everything (keyboard shortcut Ctrl+A), toggle the edge-selection icon, and select any edge of the mesh making up the border of the exposed side (Figure 12.8C). Now use Select>Select Loops>Edge Loops to automatically select the entire border (Figure 12.8D), and press keyboard shortcut “F” to create a new face within the perimeter defined by this border (Figure 12.8E). By leaving Edit Mode and going back to Object Mode (keyboard shortcut “Tab”) one can see that this gives the illusion of a solid model (Figure 12.8F), which can now be processed for illustration as described above.

### Non-conventional uses and outreach

Crucially to the paradigm outlined in Figure 3, and even though the status-quo (bias) of academia might suggest otherwise, scientific communication does not take its only form as that of a published image in a journal, nor does creative experimentation have to falter in the constraints of the printed page. As such, while 3D models do allow the generation of conventional 2D figures as described above, they also uniquely open up the doors to a wide variety of new forms of user data-engagement which are unthinkable if only dealing with pen and paper. Notably, non-conventional communication media and illustrations tools such as Augmented Reality, Virtual Reality, and 3D printing (see **Glossary**, and Figure 13), all require at their source the availability of 3D models of the objects to be brought to life (usually in either .obj or .stl format). Up to now, and in the absence of readily available compatible models, the use of these technologies to communicate mouse embryology thus required the sculpting of each of these models from scratch in software like Blender, or the generation of appropriate microscopy volume data in the lab. These requirements are unrealistic for many, either in terms of skills or opportunity, and especially since communicators and data generators might not always be the same person. Indeed examples where such approaches have been deployed are few. Yet because of their novelty, and because they engage different senses in addition than just sight, these communication strategies hold extraordinary didactic potential, whether within academia or for outreach.

**Fig. 13.**
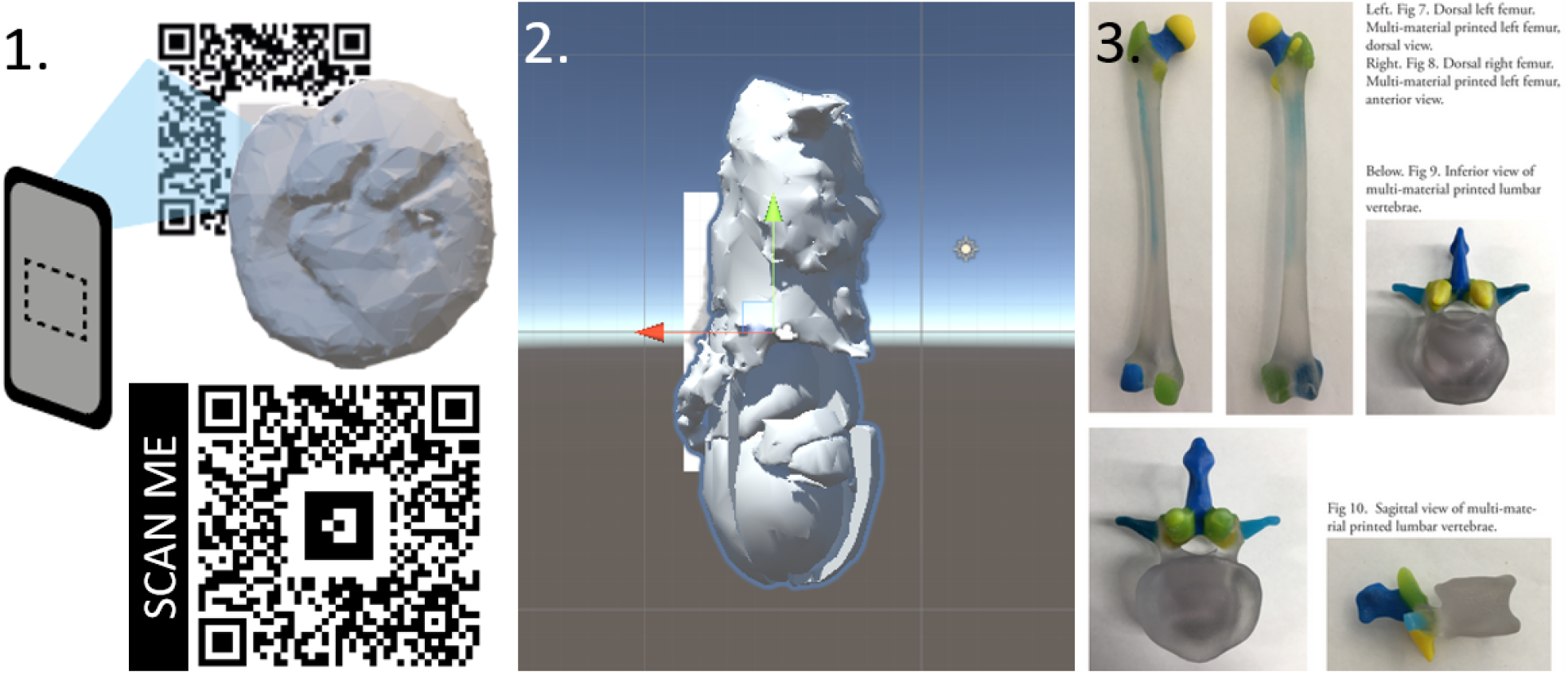
Non-conventional uses of 3D models for communication. 1. Augmented Reality to bring to life posters and papers. 2. Example of Virtual Reality setup (Unity software) 3. 3D printed models from Inoue, Leevy 2019 (https://vizbi.org/Posters/2019/B02)

The goal of this section is not to provide a detailed guide on how to deploy embryo models in virtual reality, augmented reality, or 3D printing, but just to highlight that this is possible with free and open source software. Hopefully, the embryonic models provided here can at least jump start experimentation in these avenues by interested parties. Examples of potential applications include the projection of rotating models out of posters/papers to allow live user exploration of the specific embryonic stages discussed (Figure 13.1), deployment of mouse embryos in virtual reality environments as to captivate unfamiliar but interested audiences for outreach (Figure 13.2), and multimaterial 3D printing of replica models for didactic purposes (Figure 13.3; bones in this example from https://vizbi.org/Posters/2019/B02). While a variety of different solution exist for each of these applications, augmented reality apps can be built for example with a combination of Unity and Vuforia, or the web-based AR.js. A basic script and setup behind the AR triggered by Figure 13.1 is provided at: https://github.com/StefanoVianello/Augmented_Reality. Virtual reality scenes can similarly be created by importing models in Unity and coupling this to a e.g. Google Cardboard interface; and 3D printers only require the user to supply one of the models available here. 3D printers are increasingly common in universities or in so-called community makerspaces, and we recommend considering printers with a dual nozzle setup to allow multimaterial fabrication.

## Conclusion

By converting EMAP models to a more versatile format, and by sharing them alongside models of earlier embryonic stages as ready-to-use .blend files, I hope to instill new life in such an important resource and to make its models even more accessible to the community. Undoubtedly, many researchers in the field of mouse developmental biology are in possession of 3Dmodels of various structures and embryonic stages, or generate these models as part of their research. I hope this guide can serve as a soft introduction to software (such as Blender) that allows exploration and use of these models for illustration, software that is otherwise associated with a steep learning curve and might appear overwhelming in its variety of functionalities. Hopefully, the considerations made here will also serve as an encouragement to authors to make future 3D models models they might generate as accessible as possible to the public, as their value transcend that of publication. For scientists, creators, communicators, teachers, and artists reading this, I hope these models can act as fertile substrate for creative exploration, and ultimately encourage new paradigms through which to live, communicate, and teach Developmental Biology.

## Methods

### Data availability

All data has been deposited on Zenodo (http://doi.org/10.5281/zenodo.4284380). These include: i) individual .stl models of embryonic subcomponents for each developmental stage (e.g. epiblast, visceral endoderm; 1 folder per embryonic stage); ii) ready-made .blend files where such components have been reassembled as a full embryo model, and in a scene with pre-prepared light sources and aligned camera (1 file per embryonic stage); iii) a .blend file with all embryonic models provided arranged in a temporal lineup; iv) a .blend file with a “starter-pack” of pre-made materials to use for easy illustration.

### Processing of EMAP models

EMAP models of each Theiler Stage (TS07 to TS14) were downloaded from the EMAP website (https://www.emouseatlas.org/emap/ema/home.php; also available at https://datashare.is.ed.ac.uk/handle/10283/2805) as individual .zip files. These files contain .vtk and .wlz models of i) the entire embryo and of ii) each individual embryonic subcomponent. Each folder was then decompressed (extracted), .wlz files were removed, and .vtk files were renamed according to the key provided in Supplementary Data Table. These files were then converted into .stl files through the script provided at https://doi.org/10.5281/zenodo.4284367 (Note, an alternative conversion script is also available at https://github.com/matech/PyWoolzScripts/blob/master/WlzDomainToVTKSurf.py, from Bill Hill, EMAP team). The resulting objects were finally imported into Blender 2.80, and reassembled to form a combined model of the entire conceptus for each Theiler Stage. Models corresponding to Theiler stages 01 to 06 are not available from the EMAP website and were custom made in Blender. See dedicated Methods section for more details on how these were made.

### Blender assembly of full embryo models

For each Theiler stage (TS07 to TS14), individual .stl components were imported into Blender 2.80 (File>Import>stl), shaded smooth (Object>Shade Smooth), scaled (scale: TS12 TS14 0.2. TS07-11 0.09). Origins were set to the centre of mass of each volume (Object>Set Origin>Origin to centre of mass (volume)), and an “empty” was created (Add > New > Empty) and parented to all imported pieces (control+P > Parent to Object). The whole conceptus was then assigned to a dedicated collection (group of objects, Object>Collection>Move to collection > New Collection), moved to [0,0,0], and rotated 180° with respect to its y-axis as to align the top of the model to the positive z-axis. Models were then rotated with respect to their z axis to align their posterior to the positive y axis (right side of the viewer). Finally, models were all assigned the material “base_material_white”, Color Management was kept as default (Filmic), Exposure 3, Gamma 0.75. “Film” was set as “Transparent” so that renders are created as .png by default. Note about scale: all models provided inherited size parameters from the files downloaded from EMAP. This means that TS07 to TS11 are all in scale with respect to each other and proportional to each other because they were all here rescaled by the same factor. Other models are provided unscaled and may need manual adjustment based on user needs (realism vs illustrative needs).

### Material preparation

A set of pre-made materials is available for download as to avoid having to create them from scratch. Base and Neutral materials (see Figure 5) were constructed from a single “Principled BSDF” node set on the appropriate base color (e.g. lightblue for the “neutral_lightblue” material), all other settings left to default (GGX, Christensen-Burley, Subsurface: 0, Subsurface Radius: 1-0.2-0.1, Subsurface Color: E7E7E7, Metallic: 0, Specular: 0.5, Specular Tint: 0, Roughness: 0.5, Anisotropic: 0, Anisotropic Rotation: 0, Sheen: 0, Sheen Tint: 0.5, Clearcoat: 0, Clearcoat Roughness: 0.3, IOR: 1.450, Transmission: 0, Transmission Roughness: 0, Emission: 000000, Alpha: 1). Glossy materials will give a wet-like appearance to the model, and were made just as Neutral materials but with Roughness: 0, and Clearcoat: 1. The “cells_yellow” material was created by mixing (“Mix Shader”) two “Principled BSDF” nodes with the default settings listed above for Base and Neutral Materials, with two different shades of yellow as Base Color. The “Normal” imput of the node with the lighter shade was plugged to a “Bump” node (Invert: unchecked, Strength: 1, Distance: 1) whose “Height” input was linked to the “Color” output of a “Voronoi Texture” node (Cells, Distance, Closest, Scale: 9.6). The “Fac” output of this same node was linked to a “ColorRamp” node, whose “Color” output was linked to the “Fac” input of the Mix Shader mixing the two original Principled BSDF nodes. The “cells_intercalating” material was created with the same basic setup as “cells_yellow”, but a “Musgrave Texture” node was used instead of a “Voronoi Texture” (fBM, Scale: 18.2, Detail: 4.6, Dimension: 1.614, Lacunarity: 10.5, Offse: 0.5, Gain: 0). Altering the Scale value of this node will allow to change the size of the cells as a function of the size of the tissue of interest. The “Bump” node had values= Invert: unchecked, Strength: 1, Distance: 0.2. The “transparent” material was created by mixing (“Mix Shader”) a “Principled BSDF” node set as for Base and Neutral Materials (Base Color: FFFFFF) and a “Transparent BSDF” node (Color: FFFFFF). The “Fac” value of the Mix Shader was set at 0.225 (for the transparent input).

### Preparation of pre-implantation models

Models for the 1-, 2-, and 4-cell stages (TS01.blend, TS02-2cell.blend, TS02-4cell.blend) were created from simple sphere shapes (Add>Mesh>UV Sphere). Models for the 8-cell stage, morula, early blastocyst, late blastocyst, and implanting blastocyst stages (TS03-8cell.blend, TS03-16cell.blend, TS04.blend, TS05.blend, TS06.blend) were created by using the Particle Emitter function in Blender. Briefly, an object of the desired shape (a sphere for most models; a sculpted “implanting” mesh for TS06.blend) is added (Add>Mesh>UV Sphere) and selected as an “emitter” object (“Particles” properties tab). Another object (the 1-cell model) is imported (File>Append) and selected as the “emitted” object. If the particle settings of the emitter object is set as “Hair”, running the simulation (keyboard shortcut: spacebar) will lead to individual cells emerging out of (and thus covering) the faces of the object. The entire system was then saved as a single model by applying the particle modifier (”Convert”) of the emitter object. The entire setup used to create each of the models is included in the .blend files provided, for reference and hidden from view in a dedicated collection called “Emitter_system”. The open versions of the early and mid-blastocyst models (TS04_HALF.blend, TS05_HALF.blend) were generated as above, but the emitter object used was a hollowed out hemisphere for the surface that will be covered by trophectoderm cells, and a separate shape for the surface that will be covered by inner cell mass cells (TS04_HALF.blend). For the latter particle system, the “emitted” object was a collection of two different cell objects: a sphere to represent epiblast cells, and another sphere to represent primitive endoderm cells. To instead avoiding having intermingled cells in TS05.blend, two separate inner emitters were used (each producing one type of cell).

## Supporting information

EMAP_key

Glossary

## ACKNOWLEDGEMENTS

This work would have not been possible if it were not for the existence of the Edinburgh Mouse Atlas Project. I am therefore indebted to the entire EMAP team (https://www.emouseatlas.org/emap/about/people.html) and would like to specifically thank Chris Armit for its support and encouragement at early and late stages of this project, and for his feedback on the manuscript. The guide and resources presented here are indeed intended to give new lives to EMAP models, and make them even more accessible to the community.Thanks also go to the Stack Exchange community (and specifically, user Normanius) for the code to convert volumes into .stl. I would like to thank André Dias (Instituto Gulbenkian de Ciência) for his support, correspondence, comments on the final versions of the manuscript, and for being so generous with his own 3D models. I would finally like to thank Francesca Vianello for introducing me to ARjs and for essentially setting up the entire ARjs Augmented Reality experience deployed here.

## Notes

https://zenodo.org/record/4284380

https://github.com/StefanoVianello/Augmented_Reality

https://zenodo.org/record/4284368

### Competing Interest Statement

The authors have declared no competing interest.

## Bibliography

Armit, C., Richardson, L., Venkataraman, S., Graham, L., Burton, N., Hill, B., Yang, Y., & Baldock, R. A. (2017). eMouseAtlas: An atlas-based resource for understanding mammalian embryogenesis. Developmental Biology, 423(1), 1–11. URL https://doi.org/10.1016%2Fj.ydbio.2017.01.023

Arnold, S. J., & Robertson, E. J. (2009). Making a commitment: cell lineage allocation and axis patterning in the early mouse embryo. Nature Reviews Molecular Cell Biology, 10(2), 91–103. URL https://doi.org/10.1038%2Fnrm2618

Arora, R., Fries, A., Oelerich, K., Marchuk, K., Sabeur, K., Giudice, L. C., & Laird, D. J. (2016). Insights from imaging the implanting embryo and the uterine environment in three dimensions. Development, 143(24), 4749–4754. URL https://doi.org/10.1242%2Fdev.144386

Chadwick (1978). Some aspects of the development of geological thinking. Journal of Geology Teaching, 3:142–8..

Dias, A., Lozovska, A., Wymeersch, F. J., Nóvoa, A., Binagui-Casas, A., Sobral, D., Martins, G. G., Wilson, V., & Mallo, M. (2020). A TgfRI/Snai1-dependent developmental module at the core of vertebrate axial elongation. URL https://doi.org/10.1101%2F2020.03.09.983809

Hardin, J. (2008). The Missing Dimension in Developmental Biology Education. CBE—Life Sciences Education, 7 (1), 13–16. URL https://doi.org/10.1187%2Flse.7.1.cbe13

Hashimoto, K., & Nakatsuji, N. (1989). Formation of the Primitive Streak and Mesoderm Cells in Mouse Embryos-Detailed Scanning Electron Microscopical Study. (primitive streak/cell migration/extracellular matrix/mouse gastrulation/scanning electron microscopy). Development Growth and Differentiation, 31(3), 209–218. URL https://doi.org/10.1111%2Fj.1440-169x.1989.00209.x

Ivanovitch, K., Temiño, S., & Torres, M. (2017). Live imaging of heart tube development in mouse reveals alternating phases of cardiac differentiation and morphogenesis. eLife, 6. URL https://doi.org/10.7554%2Felife.30668

Kali, Y., & Nir, O. (1996). Spatial abilities of high-school students in the perception of geologic structures. Journal of Research in Science Teaching, 33.

Lewis, S. L., & Tam, P. P. (2006). Definitive endoderm of the mouse embryo: Formation cell fates, and morphogenetic function. Developmental Dynamics, 235(9), 2315–2329. URL https://doi.org/10.1002%2Fdvdy.20846

McDole, K., Guignard, L., Amat, F., Berger, A., Malandain, G., Royer, L. A., Turaga, S. C., Branson, K., & Keller, P. J. (2018). In Toto Imaging and Reconstruction of Post-Implantation Mouse Development at the Single-Cell Level. Cell, 175(3), 859–876.e33. URL https://doi.org/10.1016%2Fj.cell.2018.09.031

Milner-Bolotin, M., & Nashon, S. M. (2011). The essence of student visual–spatial literacy and higher order thinking skills in undergraduate biology. Protoplasma, 249(S1), 25–30. URL https://doi.org/10.1007%2Fs00709-011-0346-6

Nahaboo, W., & Migeotte, I. (2018). Cleavage and Gastrulation in the Mouse Embryo. eLS. Accessed on Sat, April 11, 2020. URL https://onlinelibrary.wiley.com/doi/10.1002/9780470015902.a0001068.pub3

Nrc, N. R. C. (2012). Discipline-Based Education Research. National Academies Press. URL https://doi.org/10.17226%2F13362

Pijuan-Sala, B., Griffiths, J. A., Guibentif, C., Hiscock, T. W., Jawaid, W., Calero-Nieto, F. J., Mulas, C., Ibarra-Soria, X., Tyser, R. C. V., Ho, D. L. L., Reik, W., Srinivas, S., Simons, B. D., Nichols, J., Marioni, J. C., & Göttgens, B. (2019). A single-cell molecular map of mouse gastrulation and early organogenesis. Nature, 566(7745), 490–495. URL https://doi.org/10.1038%2Fs41586-019-0933-9

Richardson, L., Venkataraman, S., Stevenson, P., Yang, Y., Moss, J., Graham, L., Burton, N., Hill, B., Rao, J., Baldock, R. A., & Armit, C. (2013). EMAGE mouse embryo spatial gene expression database: 2014 update. Nucleic Acids Research, 42(D1), D835–D844. URL https://doi.org/10.1093%2Fnar%2Fgkt1155

Rivera-Pérez, J. A., & Hadjantonakis, A.-K. (2014). The Dynamics of Morphogenesis in the Early Mouse Embryo. Cold Spring Harbor Perspectives in Biology, 7 (11), a015867. URL https://doi.org/10.1101%2Fcshperspect.a015867

Samal, P., Maurer, P., Blitterswijk, C., Truckenmüller, R., & Giselbrecht, S. (2020). A New Microengineered Platform for 4D Tracking of Single Cells in a Stem-Cell-Based In Vitro Morphogenesis Model. Advanced Materials, (p. 1907966). URL https://doi.org/10.1002%2Fadma.201907966

Saykali, B., Mathiah, N., Nahaboo, W., Racu, M.-L., Hammou, L., Defrance, M., & Migeotte, I. (2019). Distinct mesoderm migration phenotypes in extra-embryonic and embryonic regions of the early mouse embryo. eLife, 8. URL https://doi.org/10.7554%2Felife.42434

Snow, M. H. L. (1977). Gastrulation in the mouse: Growth and regionalization of the epiblast. Development, 42: 293–303. Accessed on Sat, April 11, 2020. URL https://dev.biologists.org/content/42/1/293

Titus, S., & Horsman, E. (2009). Characterizing and Improving Spatial Visualization Skills. Journal of Geoscience Education, 57 (4), 242–254. URL https://doi.org/10.5408%2F1.3559671

Viotti, M., Foley, A. C., & Hadjantonakis, A.-K. (2014). Gutsy moves in mice: cellular and molecular dynamics of endoderm morphogenesis. Philosophical Transactions of the Royal Society B: Biological Sciences, 369(1657), 20130547. URL https://doi.org/10.1098%2Frstb.2013.0547

